## Supplemental Materials for "Developmental conversion of thymocyte-attracting cells into self-antigen-displaying cells in embryonic thymus medulla epithelium"

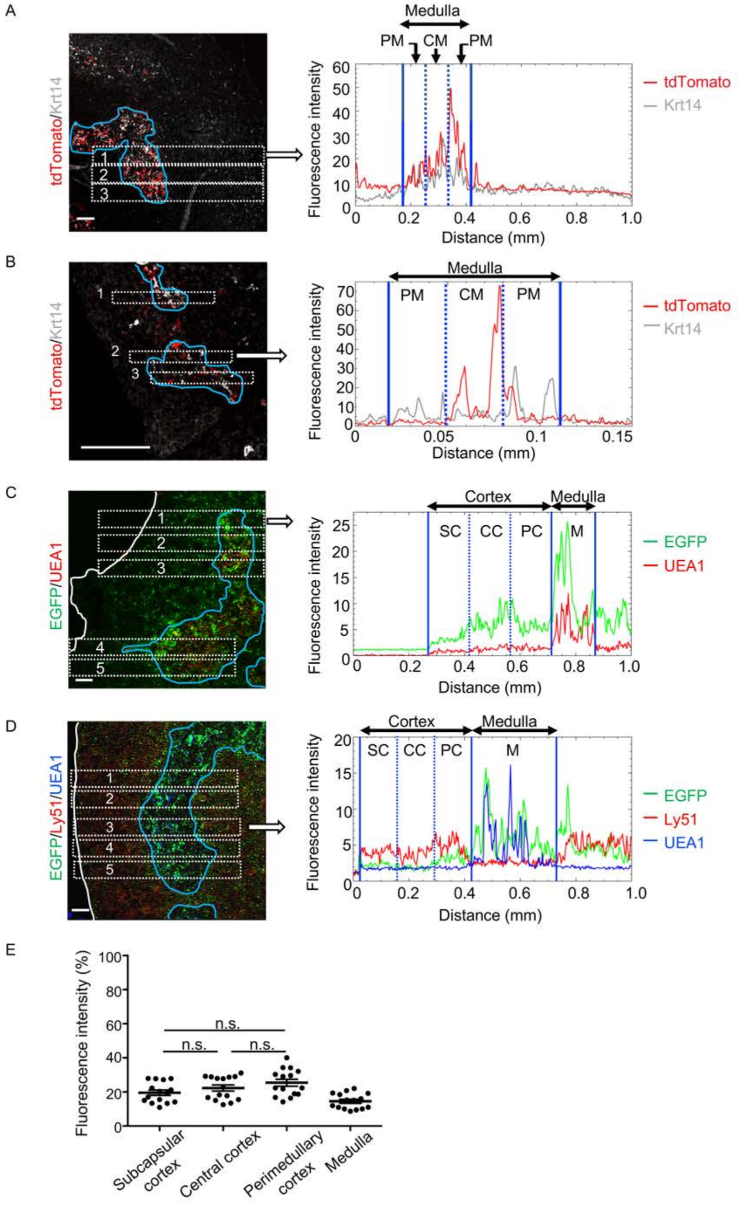


**Figure S1. Distribution of *Ccl21a*-expressing cells in the thymus.**

(**A**) A replicate of thymus section analysis for postnatal *Ccl21a*$^{tdTomato/+}$ mice as in Fig. 1E.

(**B**) A replicate of thymus section analysis for E15 *Ccl21a*$^{tdTomato/+}$ mice as in Fig. 2F.

(**C**) A replicate of thymus section analysis for postnatal *Ccl21a*-Cre x CAG-loxP- EGFP mice as in Fig. 4D.

(**D**) Distribution of Ly51 in the thymus. Left panel shows representative fluorescence signals for Ly51 (red) and EGFP (green) in the thymus sections from *Ccl21a*-Cre x CAG-loxP-EGFP mice. Right panel shows fluorescence intensity profiles of Ly51 (red), EGFP (green), and UEA1 (blue) signals within the regions of interest (ROI) defined by dashed rectangles in left panel. Cortical regions were equally divided into three areas as in Fig. 4D.

(**E**) Ly51 signal intensity (means and SEMs, n=3) in indicated areas (**D**) were calculated in comparison with total Ly51 signal intensity within the ROI. n.s., not significant.

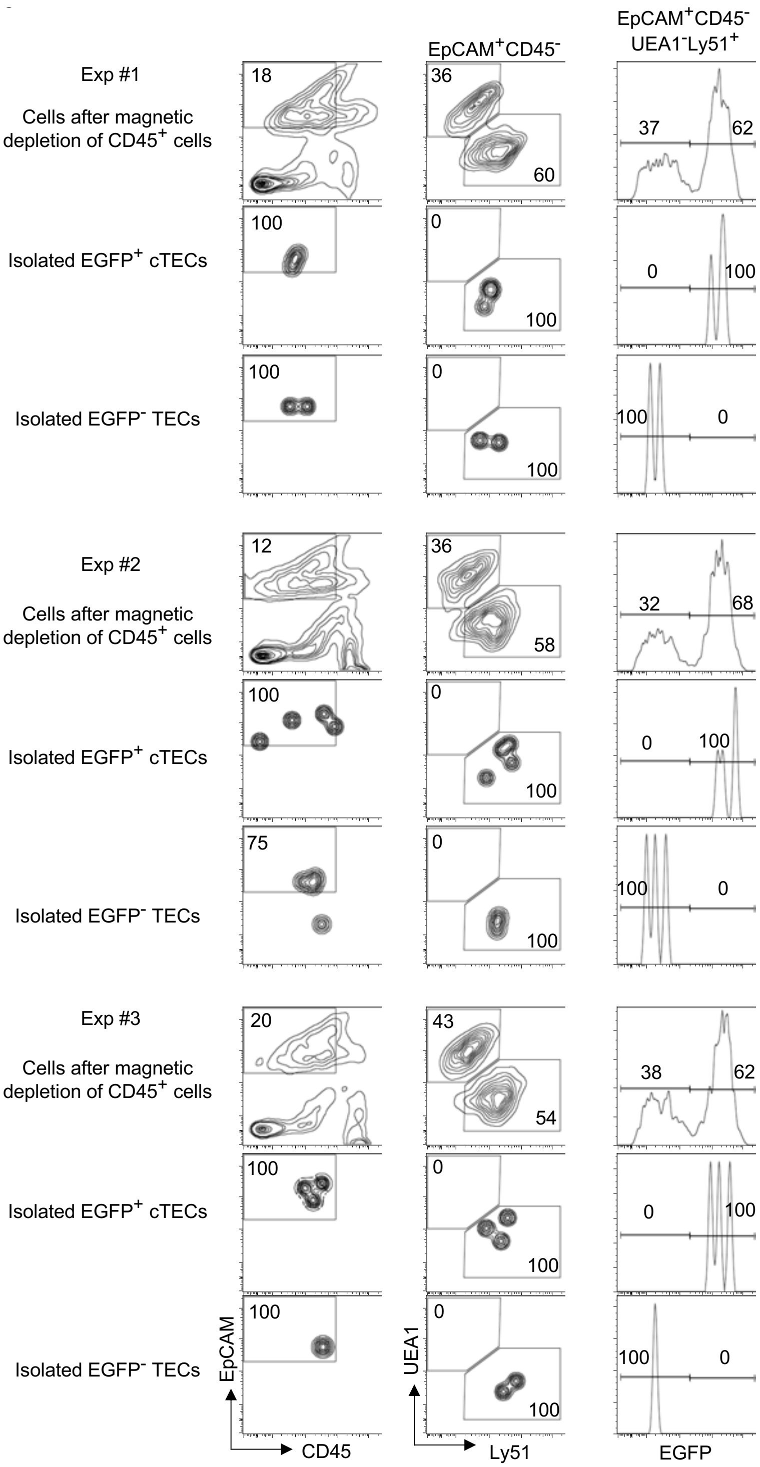


**Figure S2. Purity of isolated EGFP$^+$ and EGFP$^-$ cTECs for transcriptomic analyses.**
Flow cytometric analysis of indicated cells (n = 3) from *Ccl21a*-Cre x CAG-loxP- EGFP mice at 2-week-old. Numbers indicate frequency of cells within indicated areas.

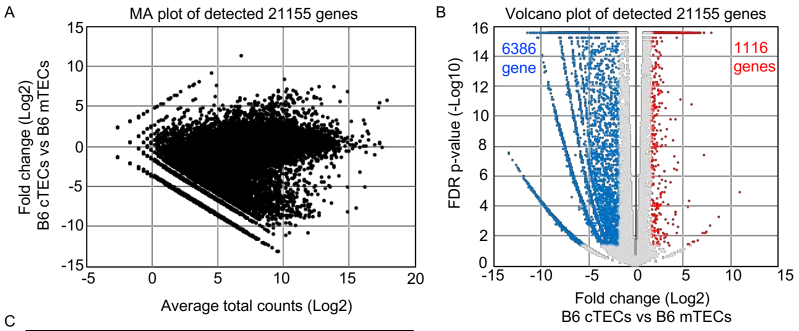


C

| | Average RPKM value | |
|---|---|---|
| Name | EGFP$^{+}$ cTEC | EGFP$^{-}$cTEC |
| Dll4 | 23.2 | 20.8 |
| Il7 | 63.0 | 66.3 |
| Cxcl12 | 110.4 | 113.8 |
| Psmb11 | 517.2 | 505.9 |
| Ctsl | 2641.4 | 2429.9 |
| Prss16 | 1670.0 | 1543.1 |
| CD83 | 441.8 | 492.9 |
| H2-K1 | 382.5 | 328.2 |
| H2-D1 | 326.5 | 301.6 |
| H2-Ab1 | 6020.6 | 5627.1 |

D

| | EGFP$^{+}$ cTEC vs EGFP$^{-}$ cTEC | | Average RPKM value | | B6 mTEC vs B6 cTEC | | Average RPKM value | |
|---|---|---|---|---|---|---|---|---|
| Name | Fold change (Log2) | FDR p-value (-Log10) | EGFP$^{+}$ cTEC | EGFP$^{-}$ cTEC | Fold change (Log2) | FDR p-value (-Log10) | B6 mTEC | B6 cTEC |
| Ccl21a | 4.0 | 9.0 | 85.9 | 5.0 | 3.9 | 16.0 | 2211.2 | 135.9 |
| Sprr2a2 | 3.8 | 9.0 | 101.3 | 7.1 | 2.3 | 16.0 | 402.4 | 76.6 |
| Sprr2a3 | 3.6 | 9.0 | 6.0 | 0.5 | 3.3 | 16.0 | 29.0 | 2.8 |
| Hbb-bh1 | 4.6 | 9.0 | 17.3 | 0.7 | 7.8 | 16.0 | 24.4 | 0.1 |
| Reg3g | 3.8 | 8.8 | 41.5 | 3.0 | 8.0 | 16.0 | 6628.2 | 24.8 |
| Sprr2a1 | 2.9 | 6.1 | 1.3 | 0.2 | 2.3 | 16.0 | 6.0 | 1.1 |
| Pth | -7.7 | 5.5 | 0.1 | 13.0 | 7.6 | 16.0 | 25.1 | 0.1 |
| Trf | 3.1 | 4.4 | 2.5 | 0.3 | 5.6 | 16.0 | 28.3 | 0.6 |
| Cxcl17 | 5.0 | 3.4 | 2.1 | 0.1 | 6.0 | 16.0 | 35.3 | 0.5 |
| Tff1 | 6.4 | 3.1 | 2.9 | 0.0 | 8.3 | 16.0 | 167.3 | 0.5 |
| Itln1 | 3.8 | 3.0 | 4.2 | 0.3 | 9.4 | 16.0 | 64.3 | 0.1 |
| Spink1 | 6.7 | 2.9 | 3.5 | 0.0 | 8.9 | 16.0 | 829.3 | 1.7 |
| Gal | 4.9 | 2.3 | 5.2 | 0.2 | 7.5 | 16.0 | 258.7 | 1.3 |
| Hbb-bt | 1.8 | 2.2 | 29.4 | 8.0 | 1.7 | 11.4 | 50.3 | 14.2 |
| Nupr1 | 1.7 | 1.3 | 4.2 | 1.2 | 5.6 | 16.0 | 345.7 | 6.7 |

**Figure S3. Transcriptomic profiles of cTECs, mTECs, EGFP$^{+}$ cTECs and EGFP$^{-}$ cTECs.**
(**A**) MA plot of 21,155 genes detected in RNA-sequencing analysis of cTECs and mTECs isolated from B6 mice[51]. Detected genes are plotted as log2 average total counts versus log2 fold changes (cTECs/mTECs).

(**B**) Volcano plot analysis of cTECs and mTECs. Detected genes are plotted as log2 fold changes (EGFP$^{+}$ cTECs/EGFP$^{-}$ cTECs) versus -log10 FDR p-values. 1,116 genes (red symbols) are more highly detected (log2 fold change > 1.7, FDR p-value < 0.05) in cTECs than mTECs, whereas 6,386 gene (blue symbol) is more highly detected (log2 fold change < -1.7, FDR p-value < 0.05) in mTECs than cTECs.

(**C**) Average RPKM value of indicated genes detected in EGFP$^{+}$ and EGFP$^{-}$ cTECs from *Ccl21a*-Cre x CAG-loxP- EGFP mice.

(**D**) Log2 fold changes, -log10 FDR p-values, and average RPKM values of genes differently detected between EGFP$^{+}$ and EGFP$^{-}$ cTECs from *Ccl21a*-Cre x CAG-loxP- EGFP mice, and those values in RNA-sequencing analysis of mTECs and cTECs from B6 mice.

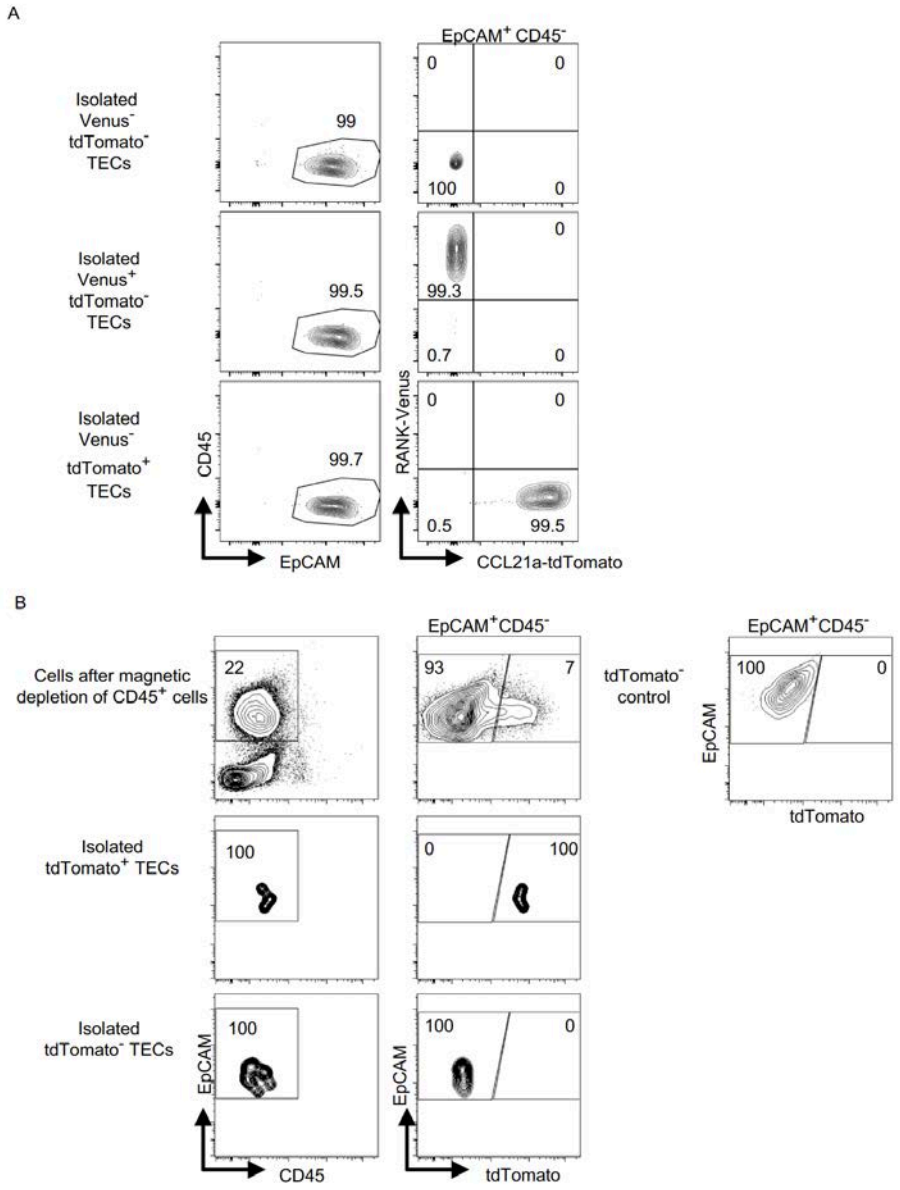


**Figure S4. Purity of isolated TECs for transcriptomic analysis and for reaggregation with relB-KO thymus stroma.**

Shown are flow cytometric profiles of indicated cells from *Ccl21a*$^{tdTomato}$ *RANK*$^{Venus}$ E17 mice (**A**) and *Ccl21a*$^{tdTomato/+}$ E17 mice (**B**). Numbers indicate frequency of cells within indicated areas.

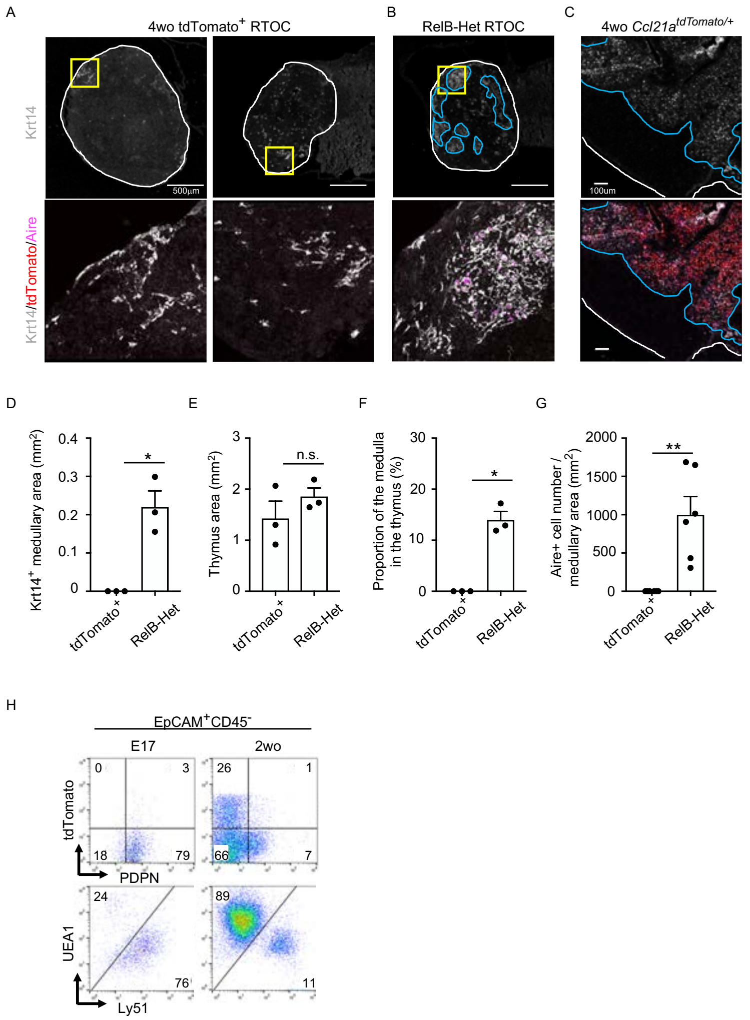


**Figure S5. Developmental potential of postnatal CCL21-expressing mTECs**

(**A, B**) Immunofluorescent staining of indicated thymus graft sections. Shown in upper panels are Krt14 expression (white). White lines indicate capsular outline of the thymus. Blue lines show Krt14$^{+}$ medullary region. Scale bar, 500µm. Bottom panels show Krt14 (white), tdTomato (red), and Aire (magenta) fluorescence signals within yellow boxes in upper panels. Grafts represent relB-KO thymic stroma reaggregated with tdTomato$^{+}$ TECs isolated from 4-week-old Ccl21a$^{tdTomato/+}$ mice (**A**) and RelB-heterozygous thymic stroma reaggregated without other TECs (**B**).

(**C**) Immunofluorescent analysis of Krt14 (white), tdTomato (red), and Aire (magenta) in thymic sections from postnatal Ccl21a$^{tdTomato/+}$ mice. Scale bar, 100µm.

(**D**) Size of Krt14$^{+}$ medullary areas (means and SEMs, n=3) in indicated thymus graft sections.

(**E**) Size of grafted thymus areas (means and SEMs, n=3) in indicated thymus graft sections.

(**F**) Proportion (means and SEMs, n=3) of Krt14$^{+}$ medullary areas in the thymus areas in indicated thymus graft sections.

(**G**) Numbers of Aire$^{+}$ mTECs per mm$^{2}$ of Krt14$^{+}$ medullary areas in indicated thymus graft sections. All images are representative data from three independent experiments. *P<0.05, **P<0.01, n.s., not significant.

(**H**) Flow cytometry profiles of tdTomato and podoplanin (PDPN) (top) and UEA1 and Ly51 (bottom) in EpCAM$^{+}$CD45$^{-}$ TECs from Ccl21a$^{tdTomato/+}$ mice at E17 and 2 weeks old. Numbers indicate frequency of cells within indicated areas.
